# Activity-based profiling of cullin-RING ligase networks by conformation-specific probes

**DOI:** 10.1101/2023.01.14.524048

**Authors:** Lukas T. Henneberg, Jaspal Singh, David M. Duda, Kheewoong Baek, David Yanishevski, Peter J. Murray, Matthias Mann, Sachdev S. Sidhu, Brenda Schulman

## Abstract

The cullin-RING E3 ligase (CRL) network comprises over 300 unique complexes that switch from inactive to activated conformations upon site-specific cullin modification by the ubiquitin-like protein NEDD8. Assessing cellular repertoires of activated CRL complexes is critical for understanding eukaryotic regulation. However, probes surveying networks controlled by site-specific ubiquitin-like protein modifications are lacking. We report development of a synthetic antibody recognizing the active conformation of a NEDD8-linked cullin. We established a pipeline probing cellular networks of activated CUL1-, CUL2-, CUL3- and CUL4-containing CRLs, revealing the CRL complexes responding to stimuli. Profiling several cell types showed their baseline neddylated CRL repertoires vary, prime efficiency of targeted protein degradation, and are differentially rewired across distinct primary cell activation pathways. Thus, conformation-specific probes can permit nonenzymatic activity-based profiling across a system of numerous multiprotein complexes, which in the case of neddylated CRLs reveals widespread regulation and could facilitate development of degrader drugs.

## Introduction

Post-translational modification by ubiquitin and ubiquitin-like proteins (UBLs) is a major mechanism for modulating protein functions in eukaryotes. Ubiquitin and UBLs toggle protein properties including half-lives, protein-protein interactions, and enzyme activities^1^. These effects are implemented by ubiquitin and UBL-binding machineries distinguishing unmodified and modified targets^2^. Some modifications, such as polyubiquitin chains with distinct linkages, are directly recognized by downstream effectors including the proteasome and Cdc48/p97. Accordingly, functions of ubiquitin chains types are amenable to probing based on ubiquitin linkage, independent of the identities of the modified proteins^3-6^. However, in many cases, regulation by monoubiquitylation, sumoylation, and neddylation involves their site-specific linkage to particular targets. These modifications and their targets are dually recognized by downstream effectors mediating regulation^7-12^. While probes for site-specifically monoubiquitylated or sumoylated targets might be restricted to a single modified protein, neddylation is distinct in that its targets include cullins, which are the core subunits in the large family of cullin-RING E3 ubiquitin ligases (CRLs)^13-15^.

Canonical CRLs are a large family of architecturally-related modular multiprotein complexes^16-19^. CRLs are assembled around elongated, multidomain cores consisting of a cullin protein paired with a RING-domain containing RBX protein (in humans, CUL1, CUL2, CUL3, CUL4A, or CUL4B with RBX1 and CUL5 with RBX2). CRL complexes form when a cullin’s N-terminal domain binds one of its dedicated, interchangeable substrate-binding modules (SBMs)^20^. CRLs are named for their constituent cullin with the variable SBM in superscript (e.g. CRL1^FBX^ E3s contain CUL1 and an F-box protein, etc). Ultimately, the cullin’s C-terminal region together with RBX1 partner with a ubiquitin carrying enzyme - an E2 such as those of the UBE2D family or RBR E3 in the ARIH family - that harbors a catalytic cysteine transferring ubiquitin to the SBM-bound substrate. The modular CRL architecture, vast number of SBMs and diversity of ubiquitin carrying enzyme partners, gives rise to a family of hundreds of unique E3 ligase complexes with distinct functions.

Cellular homeostasis depends on tight regulation of CRL activity, consistent with the irreversible impact of much of ubiquitylation. A CRL’s ubiquitin ligase function is switched on by NEDD8 linkage to a specific site conserved across cullin C-terminal so-called “WHB” subdomains^21-25^ (Fig. 1a). Modification with NEDD8 leads to a thousandfold increase in enzymatic efficiency of substrate ubiquitylation(Baek et al., 2020). The mechanism was revealed by structures visualizing ubiquitylation by neddylated CRL1 E3s: NEDD8 interacts with its covalently-linked CUL1 WHB subdomain in a specific conformation that serves as a platform for the binding and activation of ubiquitin-carrying enzymes^11,12^. The NEDD8-CUL1 WHB subdomain unit is flexibly tethered to the rest of a CRL1 complex, and oriented differently to position UBE2D-family E2s or ARIH1 adjacent to SBM-bound substrates. WHB domains from CUL1, CUL2, CUL3, and CUL4 are homologous, suggesting they form structurally similar complexes with covalently-linked NEDD8^26,27^. Indeed, mutational data confirmed the importance of the NEDD8-CUL4 interface in degrader-drug-induced ubiquitylation^11^. However, the sequence of CUL5 lacks key features of the interface with NEDD8, and is activated through a distinct structural mechanism^28^.

**Fig. 1:**
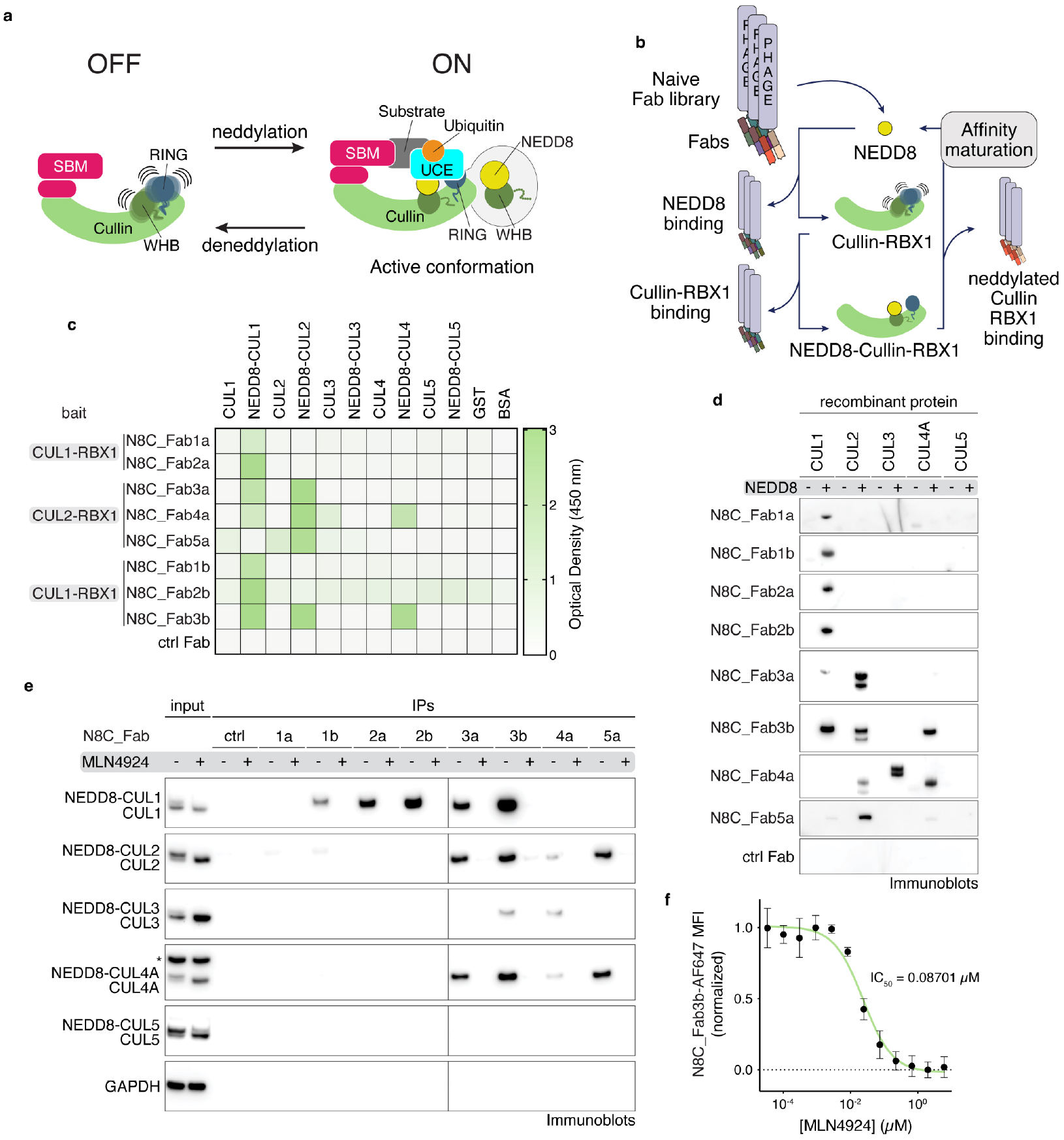
A suite of synthetic antibody fragments (Fabs) specific for NEDD8-modified cullin-RING ligases (CRLs). **a**, CRLs are switched on by site-specific NEDD8 linkage to the cullin’s WHB domain. Neddylation promotes the active conformation required for ubiquitylation to be assumed (SBM = substrate binding module, UCE = ubiquitin carrying enzyme). **b**, Strategy to select Fabs specifically binding to the NEDD8-modifed, and not unmodified, CRLs. Selections were performed with neddylated C-terminal regions of CUL1 or CUL2 bound to RBX1. **c**, Binding specificity of Fabs developed with selection strategy seen in (b) against non-neddylated and neddylated CUL1-5, GST and BSA as determined by ELISA. Baits used for selection of specific Fabs are indicated. **d**, Immunoblot using indicated Fabs of the suite as primary binders for recognition of indicated purified recombinant cullins modified by NEDD8 (+) or not (-). **e**, IP with indicated Fabs from K562 (human lymphoblast) cells treated with DMSO or MLN4924, followed by immunoblotting against cullins 1-5. Slower migrating forms of cullins eliminated by MLN4924 treatment are interpreted as NEDD8-modified, whereas faster-migrating forms of cullins are interpreted as unneddylated. * - band cross-reacting with anti-CUL4 antibody. **f**, Dose-response curve of MLN4924 in K562 cells as measured by flow cytometry using N8C_Fab3b fluorescently labeled with Alexa Fluor 647 as a direct read-out of cullin neddylation levels.

As neddylation determines which CRLs are switched on, this is tightly controlled^13^. Specialized E1-E2-E3 cascades catalyze NEDD8 linkage to cullins on the millisecond time scale. Meanwhile, CSN deconjugates NEDD8 in reactions operating near the diffusion-controlled limit. Neddylation is a fragile modification that is rapidly reversed unless bound to a substrate, which sterically protects CRLs from CSN-catalyzed deneddylation^29-34^. Thus, NEDD8 linkage to a cullin is a marker of a substrate-bound, active CRL.

With NEDD8 and CRLs implicated in virtually every facet of eukaryotic biology, it is of great interest to probe NEDD8-activated CRLs. Indeed, regulation of the CRL network by neddylation has been implicated in the cell cycle, immune signaling, DNA replication and repair, oxidative stress response, hypoxia signaling, bacterial hijacking, viral hijacking and tumorigenesis^16,18,20^. Neddylation is also important in CRL-dependent targeted protein degradation^35-38^. Furthermore, endogenous tagging of specific cullins and quantitatively surveying which SBMs are impacted by inhibiting neddylation^39-42^ led to an “adaptive exchange hypothesis” proposing that the landscape of NEDD8-activated CRLs is rewired to adapt to changes in cellular conditions. However, current methods to assess CRL repertoires require endogenous tagging^40,42^, which is laborious, may introduce artefacts, limits studies to the single tagged cell line and could prove challenging for primary cells.

In this work, we generate antigen-binding fragments (Fabs) via phage display that selectivity target neddylated cullins, with nanomolar affinities. Structural studies show that one of the generated Fabs binds neddylated CUL1 in the active conformation during ubiquitylation, while biochemical and proteomics reveal it captures neddylated CRL1, CRL2, CRL3, and CRL4 complexes with high specificity. We developed a quantitative proteomics pipeline using this activity-based probe to profile distinct active CRL complex landscapes, with multi-cullin coverage. Application to various human and murine cells revealed widespread signaling through rearrangement of complex networks in response to diverse stimuli. Overall, our probing the active conformation of specific ubiquitin-like protein-modified targets discovered widespread regulation of numerous E3 ligases that is not reflected by changes in the modification itself or in protein abundances. Moreover, we provide a robust method for probing active CRL signaling pathways and informing cellular readiness for small molecule degraders.

## Results

### A suite of synthetic antibodies specifically recognizing NEDD8-modified cullin-RING ligases

We sought to generate probes selectively binding neddylated CRLs. Thus, we established a negative-positive selection strategy to enrich for specific binders from a library of Fabs on phage^43-45^. First, the library was depleted of Fabs recognizing either a cullin-RING complex or NEDD8 without the other. Next, a neddylated cullin-RING complex was the bait for positive selection (Fig. 1b, Extended Data Fig. 1a,b). The baits were minimal complexes between RBX1 and the C-terminal regions of cullins that can be enzymatically neddylated^21^. Performing independent selections with two different cullins, CUL1 or CUL2, yielded two and three Fab sequences, respectively.

The affinities and specificities of the selected phage-displayed Fabs were assessed by enzyme-linked immunosorbent assays (ELISAs) (Extended Data Fig. 2a-c). The Fabs specifically bound neddylated cullins, showing little to no binding to unneddylated cullins, GST or BSA (Fig. 1c). Panning across the cullin family showed several Fabs were specific for the cullins used as baits in their selection. Unexpectedly, however, two Fabs selected to bind a neddylated version of CUL2-RBX1 showed broader interactions, with neddylated CUL1-RBX1, and one also with neddylated CUL4A-RBX1.

**Fig. 2:**
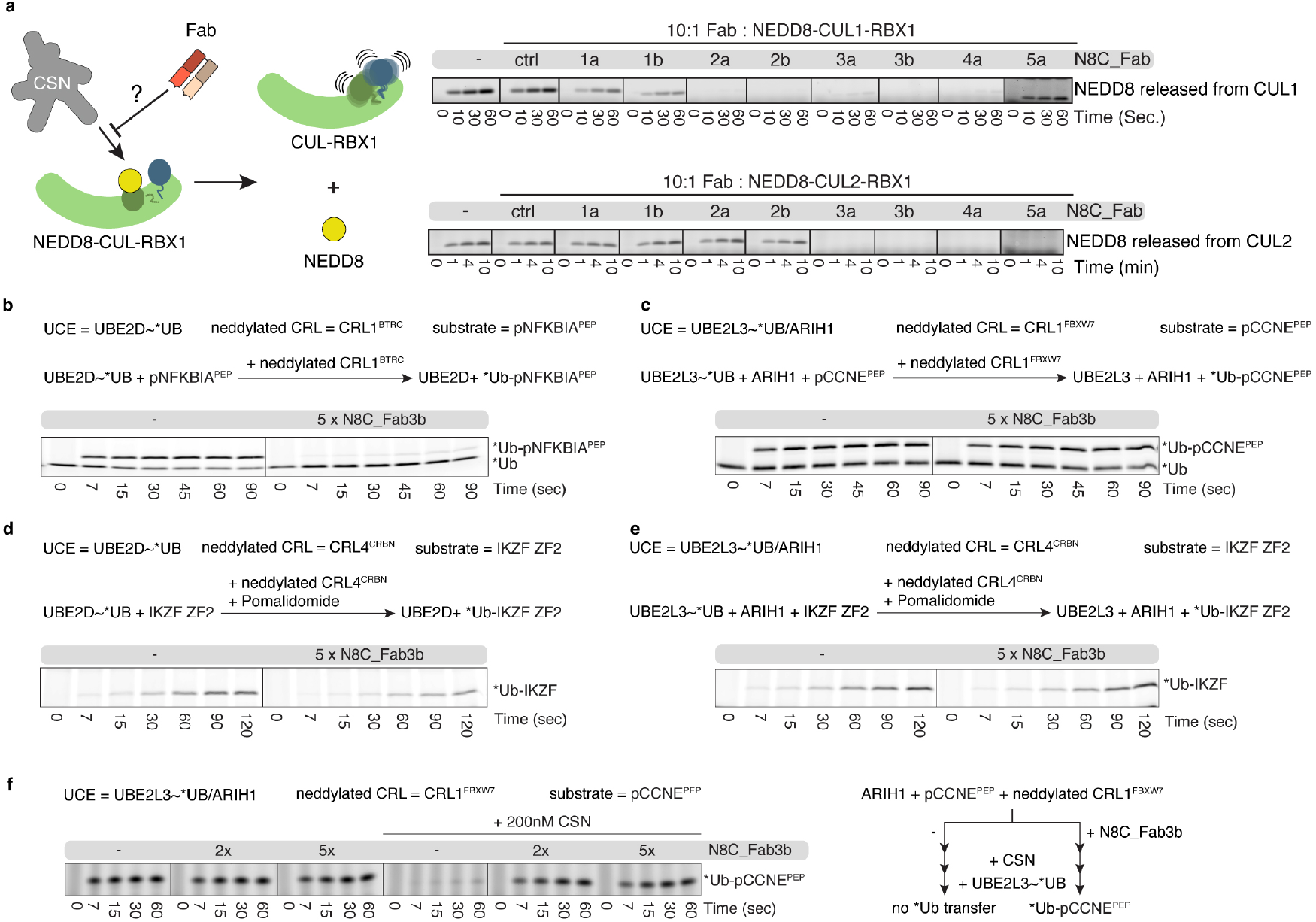
Effects of neddylated cullin targeting Fabs on CRL activities. **a**, CSN-catalyzed NEDD8 deconjugation is visualized by accumulation of free NEDD8 observed over time by SDS-PAGE. Assays test protective effects of incubating 10x molar excess of indicated Fabs with neddylated C-terminal regions of CUL1 or CUL2 complexes with RBX1. **b**, Effects of N8C_Fab3b binding on ubiquitylation efficiency of neddylated CRL1^BTRC^-UBE2D (aka CRL1^βTrCP1^-UBE2D) as determined by a qualitative substrate priming assay monitoring transfer of fluorescent ubiquitin (*Ub) to a phospho-NFKBIA (aka phospho-IκB⍺) derived substrate peptide (pNFKBIA^PEP^) by SDS-PAGE. **c**, As (b), but monitoring the transfer of *Ub by neddylated CRL1^FBXW7^-UBE2L3/ARIH1 to a phospho-CCNE (aka phospho-Cyclin E) derived substrate peptide (pCCNE^PEP^). **d**, As (b), but monitoring Pomalidomide induced transfer of *Ub by neddylated CRL4^CRBN^-UBE2D to the IKZF ZF2 substrate. **e**, As (b), but monitoring Pomalidomide induced transfer of *Ub by neddylated CRL4^CRBN^-UBE2L3/ARIH1 to the IKZF ZF2 substrate. **f**, As (c), but comparing *Ub transfer in the absence and presence of CSN with or without prior incubation with N8C_Fab3b.

To further improve affinities for CUL1, and to investigate if an orthogonal selection could extend the range of cullins recognized by a single Fab, we performed another round of selections using the neddylated CUL1 fragment bound to RBX1 as the bait. New libraries were based on the sequences of N8C_Fab1a, 2a, and 3a, with soft randomization in their complementarity-determining regions (CDR)-L3 and H3. Selections with the libraries based on N8C_Fab1a and 2a yielded new Fabs with up to 3-fold increased affinity compared to their original counterparts (Extended data Fig. 2b). The selection with the library based on the framework of N8C_Fab3a, which came from the screen against neddylated CUL2-RBX1, led to a Fab with remarkable properties by ELISA: maintaining interaction with neddylated CUL2-RBX1, 20-fold improvement in EC_50_ toward neddylated CUL1-RBX1, and emergent recognition of neddylated CUL4A-RBX1. Overall, the selections yielded eight Fabs, some specifically binding neddylated versions of either CUL1-RBX1 or CUL2-RBX1, while others recognized multiple neddylated cullin-RBX1 complexes with low nanomolar EC_50_s.

### N8C_Fabs selectively detect neddylated cullins by immunoblotting and immunoprecipitation

We tested purified versions of the Fabs for selective detection of neddylated cullins in various assays. In a western blot format, all the Fabs specifically recognized neddylated cullins. Here, cullin preferences correlated with those of the phage-displayed Fabs detected by ELISA (Fig. 1d). The trends also held when the Fabs were used to perform immunoprecipitation (IP) from K562 cell lysates, followed by immunoblotting with commercial cullin-specific antibodies. Neddylation-dependence was confirmed by the elimination of interactions from cells treated with the neddylation inhibitor MLN4924^46^ (Fig. 1e). Although neddylated CUL3-RBX1 was not detected by ELISA as interacting with any of the Fabs when displayed on phage, it was detected by purified N8C_Fab4a in a western blot and enriched by immunoprecipitation with both N8C_Fab3b and N8C_Fab4a from cell lysates.

Based on its high specificity for neddylated over non-neddylated cullins, and capacity to bind multiple cullins, we tested N8C_Fab3b for utility in flow cytometry. If the N8C_Fab3b directly detected neddylated cullins, then signal would be eliminated by treatment with MLN4924. Indeed, the dose-response curves for K562 cells showed a half-maximal inhibitory concentration (IC_50_) of ∼87 nM, in line the <100 nM previously estimated based on NEDD8 migration detected in western blots as a proxy for conjugate formation^46^.

### N8C_Fabs effects on neddylated CRL activities

Neddylation alters the binding partners and functions of cullin-RING complexes. Thus, we tested the effects of adding N8C_Fabs to activity assays. First, we explored whether any of the Fabs could protect the fragile NEDD8 mark from deconjugation by CSN. Whereas CSN rapidly catalyzed NEDD8 removal from CUL1 and CUL2 complexes with RBX1, addition of several of the N8C_Fabs to these reactions slowed deneddylation (Fig. 2a). Retention of the NEDD8 linkage largely correlated with Fab binding measured by ELISA (Extended Data Fig. 2b), with some exceptions. For example, N8C_Fab4a provided robust protection of neddylated CUL1 compared to N8C_Fab1b despite almost 3-fold higher EC_50_. A possible explanation for the differences would be if only a subset of the Fabs bind in such a way as to prevent CSN access to the NEDD8-cullin bond.

We selected N8C_Fab3b for further characterization based on its binding the broadest range of neddylated cullins, and because it maintained cullin neddylation in the presence of CSN. We tested effects on structurally-characterized ubiquitylation reactions, with an E2 (UBE2D), or E3 (ARIH1, which collaborates with the E2 UBE2L3 to ubiquitylate CRL substrates)^24^. N8C_Fab3b inhibited UBE2D- and neddylated CRL1^BTRC^-dependent (aka βTrCP1) ubiquitylation of a peptide substrate derived from the phospho-NFKBIA (aka IκB⍺) (Fig. 2b). Meanwhile, there was no effect of adding N8C_Fab3b on UBE2L3/ARIH1- and neddylated CRL1^FBXW7^-mediated ubiquitylation of a peptide substrate derived from phospho-Cyclin E (Fig. 2c). The distinct effects of N8C_Fab3b correlated with the ubiquitin carrying enzyme used in the reaction, as shown by examining pomalidomide-induced CRL4^CRBN^-mediated ubiquitylation of a peptide substrate based on the Ikaros degron (Fig. 2d, e).

The lack of effects of N8C_Fab3b on reactions with ARIH1 could be explained in two ways. N8C_Fab3b simply might not bind during ARIH1-dependent ubiquitylation. Alternatively, N8C_Fab3b might be compatible with ubiquitylation by ARIH1. This latter possibility is of interest, because cullins 1-4 and a large fraction of CUL1, CUL2, and CUL3-associated SBMs co-immunoprecipitate with ARIH1; these interactions are neddylation-dependent as they are eliminated by cell treatment with MLN4924^24^. Thus, we devised an experiment distinguishing the possibilities, based on competition between N8C_Fab3b and deneddylation, and requirement for CRL neddylation for ARIH1-mediated ubiquitylation (Fig. 2f). In accordance with its potent deneddylation activity, adding CSN at a concentration overcoming CRL substrate inhibition eliminated UBE2L3/ARIH1-mediated ubiquitylation. However, ubiquitylation activity was restored by N8C_Fab3b. Thus, N8C_Fab3b protects the neddylated CRL1 complex during the ubiquitylation reaction.

### N8C_Fab3b captures the active conformation of NEDD8-CUL1 complex

To understand the molecular basis for selective recognition, we sought to determine the structure of a N8C_Fab3b complex with a neddylated cullin. Thus, we devised a strategy to isolate enzymatically neddylated CUL1 WHB domain (Extended Data Fig. 3a-c), which co-crystallized with N8C_Fab3b, yielding a structure at 2.7 Å resolution. There are two N8C_Fab3b-neddylated CUL1^WHB^ complexes in the asymmetric unit. These superimpose with a root mean square deviation of 0.459 Å, so only one complex is described (Extended Data Table 1).

**Fig. 3:**
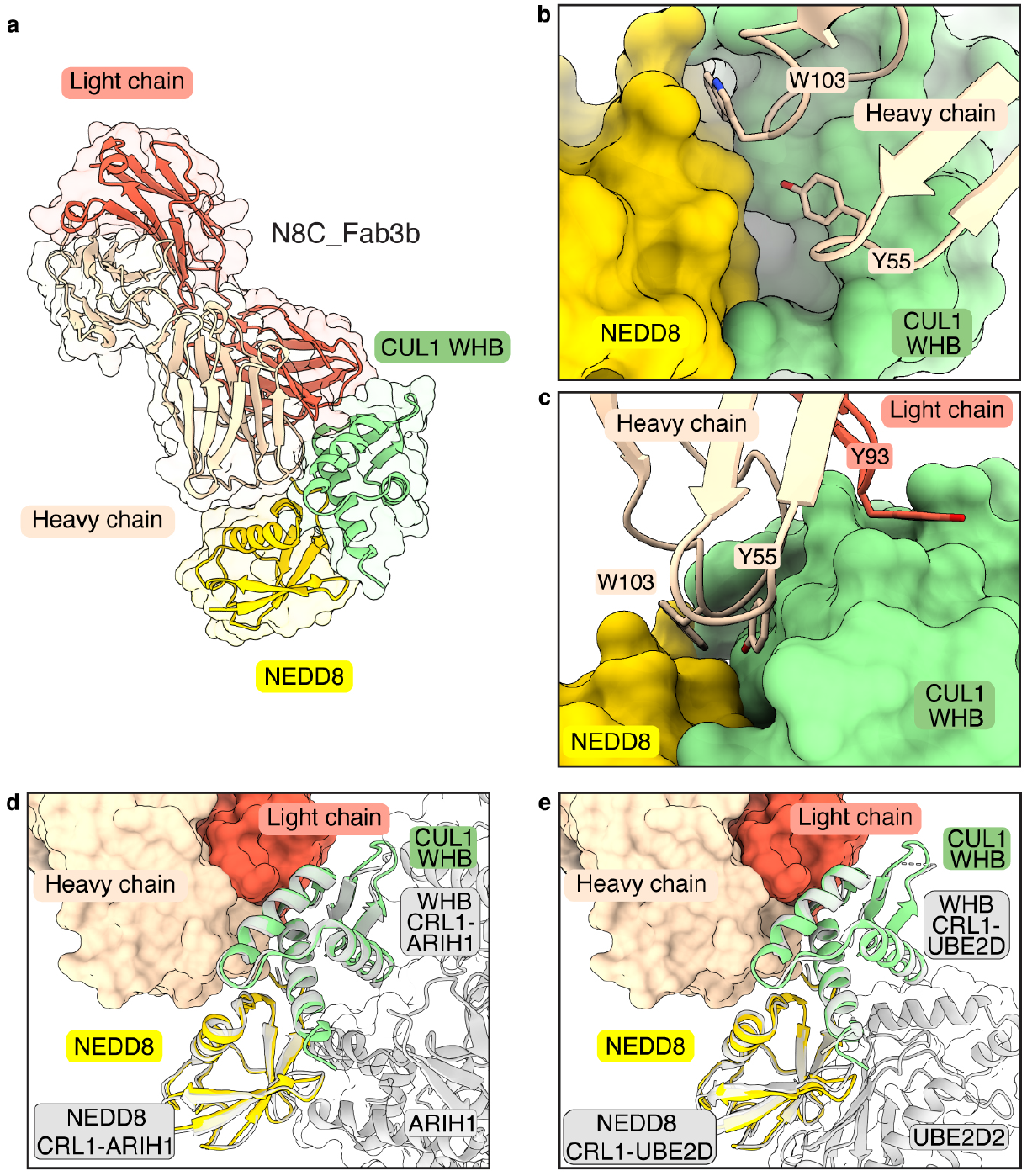
Crystal structure of N8C_Fab3b in complex with neddylated CUL1^WHB^ reveals capture of the active conformation. **a**, Crystal structure of N8C_Fab3b in complex with neddylated CUL1^WHB^ at 2.6 Å resolution. N8C_Fab3b recognizes a unique interface spanning both NEDD8 and CUL1. **b**, Focused view of the N8C_Fab3b wedge consisting of residues Y55 and W103 of the Fab heavy chain burying itself in the groove between the CUL1 WHB domain and NEDD8. **c**, The N8C_Fab3b wedge is further stabilized by Y93 hooking into the edge of the CUL1 WHB domain. **d**, Side-by-side comparison of the NEDD8-CUL1^WHB^ bound by N8C_Fab3b with the one seen in an active CRL1^FBXW7-^ UBE2L3/ARIH1 complex (7B5L) reveals N8C_Fab3b capturing the active conformation of NEDD8-CUL1^WHB^. Alignments were performed on NEDD8-CUL1^WHB^. **e**, Alignment of CUL1^WHB^-NEDD8 bound by N8C_Fab3b and the one found in the active CRL1^BTRC^-UBE2D2 complex (6TTU) further demonstrates N8C_Fab recognizing the active NEDD8-CUL1^WHB^ conformation and being compatible with ubiquitin carrying enzymes. Alignments were performed on NEDD8-CUL1^WHB^.

The structure reveals the strict requirement for cullin neddylation: N8C_Fab3b binds a unique interface spanning both NEDD8 and CUL1 (Fig. 3a). Tyr55 and Trp103 of N8C_Fab3b CDR-H2 and H3, respectively, insert into a groove between NEDD8’s so-called Ile36 patch and the CUL1 WHB domain. One edge of this groove is the isopeptide linkage between NEDD8 and CUL1, and the other is established by noncovalent NEDD8-CUL1 contacts (Fig. 3b). The complex is stabilized by multiple hydrogen bonds between the CDRs of N8C_Fab3b and either NEDD8 or CUL1 (Extended Data Fig. 4a), as well as Tyr93 of CDR-L3 clasping the edge of CUL1’s WHB domain (Fig. 3c). As such, N8C_Fab3b binds a specific arrangement of NEDD8 and its linked CUL1 WHB domain.

**Fig. 4:**
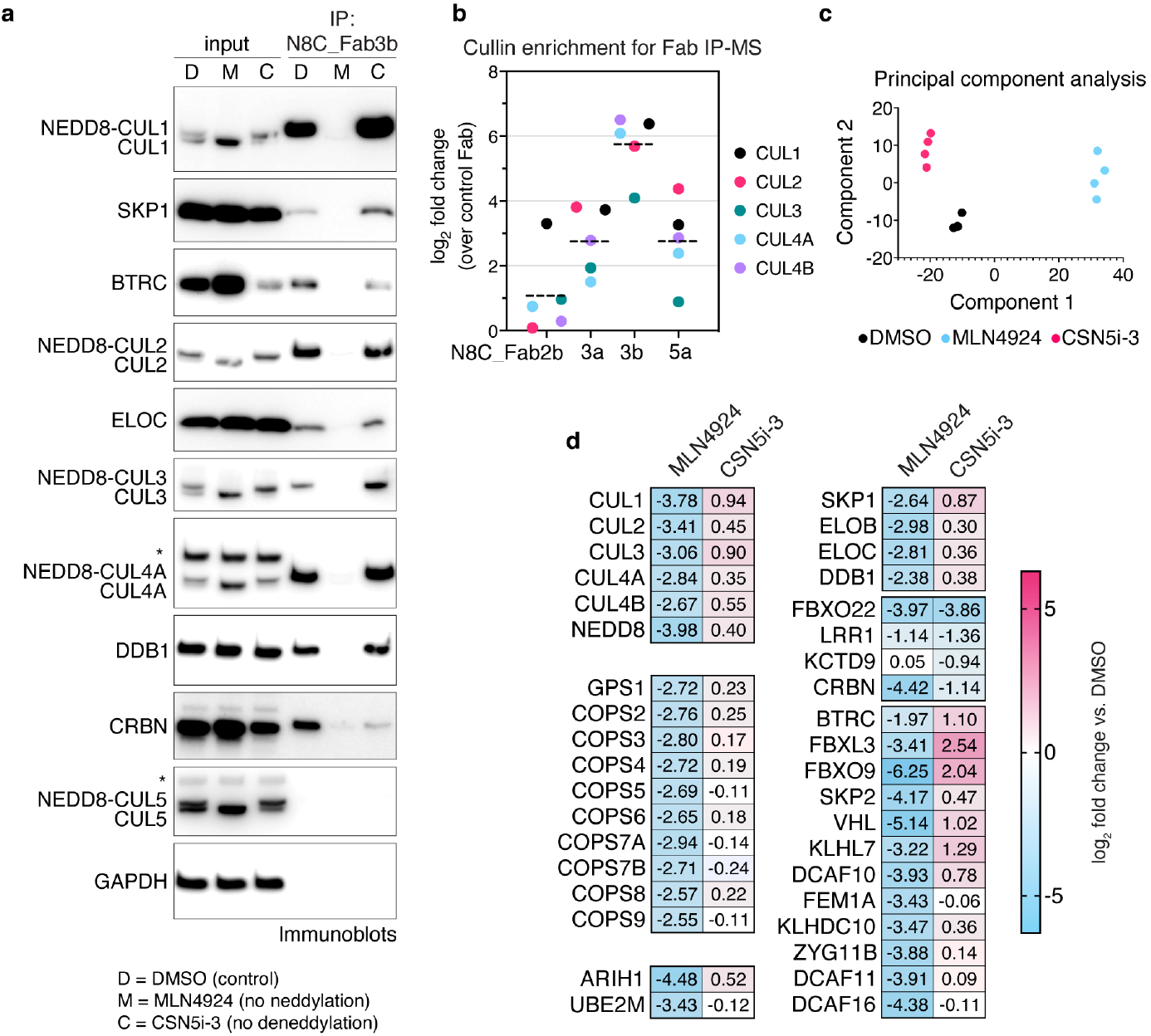
Probing the cellular repertoire of active CRLs. **a**, IPs with N8C_Fab3b from K562 (human lymphoblast) cells treated with DMSO (D), MLN4924 (M) or CSN5i-3 (C), probed for known CRL components by immunoblotting. Slower migrating forms of cullins eliminated by MLN4924 treatment are interpreted as NEDD8-modified, whereas faster-migrating forms of cullins are interpreted as unneddylated. * - band cross-reacting with anti-CUL4 or anti-CUL5 antibody. **b**, Comparison of enrichment of cullins seen in IP-MS experiments with Fabs N8C_Fab2b, 3a, 3b and 5a compared to a control Fab bearing wild type CDRs. **c**, PCA plot of IP-MS experiments with N8C_Fab3b from 293T cells treated with DMSO, MLN4924 or CSN5i-3 for 2h. **d**, Heatmap of selected proteins significantly differentially identified after either MLN4924 or CSN5i-3 treatment in (c).

Remarkably, the arrangement of the N8C_Fab3b-bound NEDD8-CUL1^WHB^ unit matches that in structures representing neddylated CRL1^BTRC^ (aka βTrCP1) ubiquitylation with the E2 UBE2D(Baek et al., 2020) and neddylated CRL1^FBXW7^ ubiquitylation with the E2/E3 UBE2L3/ARIH1(Horn-Ghetko et al., 2021) (Fig. 3d and e). Notably, the key WHB domain residues binding noncovalently to NEDD8 are conserved in cullins 1-4 (Extended Data Fig. 4b). However, CUL5’s sequence is incompatible with forming such a complex; NEDD8 and CUL5’s WHB domain adopt a different conformation in neddylated CRL5 E3s^28^ (Extended Data Fig. 4c). Thus, selectivity is determined not only by the interactions mediated directly by N8C_Fab3b, but also by the capacity for a neddylated CRL to form the active assembly between NEDD8 and a cullin’s WHB domain.

NEDD8 and CUL1’s WHB domain are thought to sample multiple conformations in a neddylated CRL1 complex: they were not visualized in prior cryo EM structures without a ubiquitin carrying enzyme, and the NEDD8-CUL1 WHB domain unit occupies different relative positions to activate UBE2D or ARIH1^11,12^. Docking N8C_Fab3b onto the prior structures of ubiquitylation complexes shows that although it would clash during ubiquitin transfer from UBE2D to an SBM-bound substrate, this Fab can readily capture an ARIH1-bound CRL complex (Extended Data Fig. 4d). This explains effects of N8C_Fab3b on the different ubiquitylation reactions (Fig. 2b-e). The structure with N8C_Fab3b also suggests NEDD8 and a cullin’s WHB domain have intrinsic propensity to bind each other in the active conformation. Together, the data show N8C_Fab3b captures the active arrangement between NEDD8 and its linked cullin domain, and reveal its potential to probe for NEDD8-activated CRLs.

### A pipeline probing cellular repertoires of neddylated CRLs

Given that N8C_Fab3b IPs could enrich NEDD8-activated CRLs, we next optimized conditions to establish a pipeline meeting several key criteria. First, we sought to identify proteins interacting specifically with neddylated cullins. This was distinguished by comparing effects of cell treatment with DMSO as control or MLN4924 that eliminates cullin neddylation^46^. We also examined effects increasing cullin neddylation, by treating cells with the CSN inhibitor CSN5i-3^47^. Second, an elaborate CRL assembly and disassembly pathway is thought to shuffle the limited cellular pool of cullin-RBX1 subcomplexes between excess SBMs in a deneddylation-dependent process. CRL disassembly is transiently paused by retention of NEDD8 on the CRLs bound to substrates. To preserve the cellular repertoire of neddylated CRLs, post-lysis cullin reshuffling must be prevented by a “N8-block”, where MLN4924 and CSN5i-3 are applied during harvesting, and included in lysis and wash buffers^42^. Performing IPs with N8C_Fab3b using an N8-block, and immunoblotting indeed showed neddylation-dependent enrichment of CRL components such as adapter proteins SKP1, ELOC and DDB1, and SBMs including BTRC (aka βTrCP1) and CRBN (Fig. 4a). To confirm that N8C_Fab3b is most suited for profiling active CRL interactors, we also performed the IPs with N8C_Fab2b, N8C_Fab3a, and N8C_Fab5a, followed by library-free data independent acquisition (DIA) mass spectrometry (MS). The Fabs all significantly enriched cullins and SBMs in a neddylation-dependent manner (Fig. 4b, Extended Data Fig. 5a). However, cullin specificities generally paralleled their in vitro binding properties (Extended Data Fig. 1c). N8C_Fab3b on average yielded 10-fold greater enrichment of known cullin-associated proteins compared to the next best Fab, so we selected it to profile cellular activated CRL-omes (Fig. 4b).

**Fig. 5:**
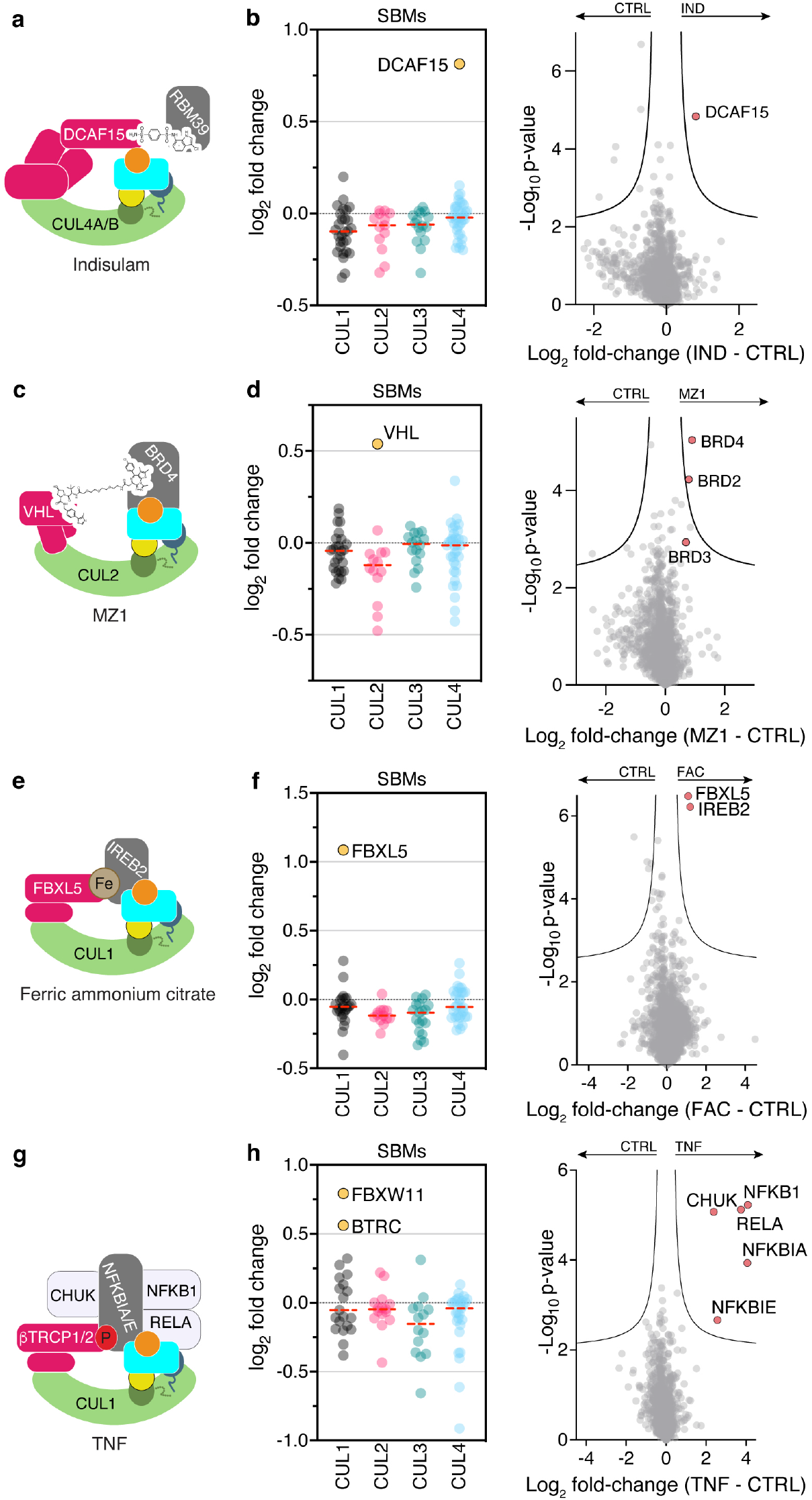
Profiling CRL complexes activated by extracellular signals. **a**, Indisulam recruits RBM39 as a neo-substrate for degradation by CRL4^DCAF15^. **b**, Shift seen for SBMs associated with cullins 1-4 in N8C_Fab3b profiling (left) and across all identified proteins after 1h treatment of 293T cells with 2 µM Indisulam (IND). **c**, MZ1 hijacks CRL2^VHL^ for degradation of BRD4. **d**, Same as (b) but after 1h treatment of 293T cells with 1 µM MZ1. **e**, CRL1^FBXL5^ degrades IREB2 in an iron dependent manner. **f**, Same as (b) but after 90 min treatment of 293T cells with 100 µM ferric ammonium citrate (FAC). **g**, TNF stimulation causes phosphorylation of NFKBIA and NFKBIE and their subsequent ubiquitylation by CRL1^BTRC^ and CRL1^FBXW11^ (aka CRL1^βTRCP1^ and CRL1^βTRCP2^). **h**, Same as (b) but after 5 min treatment of K562 cells with 25 ng/ml TNF.

We next used unbiased proteomics in DIA format to probe the interactome of N8C_FAB3b. IPs from 293T cells treated with DMSO, MLN4924 or CSN5i-3 for 2 hours readily identified interactors known to vary across the differing neddylation states imposed by the inhibitors, providing positive controls (Fig. 4c). Besides the cullins and NEDD8, the interactome included components of the COP9 signalosome, the RBX1-specific NEDD8 E2 UBE2M and ubiquitin carrying enzyme ARIH1, and adapter proteins SKP1, ELOB, ELOC and DDB1, all of which showed strong neddylation dependence (Fig. 4d). Notably, the CUL5-RBX2-specific NEDD8 E2 UBE2F and ubiquitin carrying enzyme ARIH2 were not observed, indicating that the interactors are with neddylated cullins 1-4 recognized by N8C_Fab3b.

Interestingly, two categories of neddylation-dependence were observed for SBM interactions. SBMs in one category, which includes CRBN, decrease with either MLN4924 or CSN5i-3 treatment (Fig. 4d). This behavior might be explained by autoubiquitylation-mediated degradation, as has been previously reported for CRBN(Mayor-Ruiz et al., 2019). However, the majority decrease with MLN4924, and either remain similarly bound or increase with CSN5i-3 treatments (Fig. 4d).

### Profiling CRL complexes activated by extracellular signals

The results from Figure 4d suggested that N8C_Fab3b could identify CRL complexes switching neddylation state in response to external stimuli. To explore this further, we tested three types of stimuli known to trigger neddylated CRL-dependent protein degradation, using a protocol that inhibits protein turnover.

First, we profiled responses to two types of degrader drugs. We examined a molecular glue, Indisulam, which engages neddylated CRL4^DCAF15^ to degrade RBM39^48,49^ (Fig. 5a). Profiling with N8C_Fab3b identified DCAF15 as increasing in neddylated cullin association (Extended Data Fig. 5b). Strikingly, this was the only significant change triggered by Indisulam (Fig. 5b). We next examined effects of a Proteolysis Targeting Chimera (PROTAC), MZ1, that affixes complexes between CRL2^VHL^ and Bromo- and Extra-Terminal domain (BET) family members BRD2, BRD3 and BRD4^50^ (Fig. 5c). Our active CRL profiling showed MZ1 triggers enrichment of VHL (Extended Data Fig. 5c). It also revealed BRD2, BRD3 and BRD4 as associating with a neddylated CRL in an MZ1-dependent manner (Fig. 5d). Thus, our profiling method can identify the CRL activated by a degrader drug, and in some cases targets for degradation as well. The preferential enrichment of BRD4 over BRD2 and BRD3 correlates with MZ1-targeted degradation rather than its affinity for these neo-substrates^50^, in accordance with the concept that CRL complex architecture rather than substrate binding is the driver of degradation^51,52^.

Many endogenous cellular signaling pathways depend on CRL-based responses to execute biological functions. To determine if profiling with N8C_Fab3b permits capturing such regulation, we first examined a metabolic signaling pathway where high iron levels trigger CRL1^FBXL5^-dependent degradation of iron regulatory protein 2 (IREB2)^53,54^ (Fig. 5e). Indeed, treatment of cells with ferric ammonium citrate elicited ∼2-fold relative increases only in FBXL5 and its substrate IREB2 as neddylated cullin-associated proteins (Fig. 5f, Extended data Fig. 5d). Finally, we also profiled the response to the cytokine TNF, which induces degradation of phosphorylated NFKBIA and NFKBIE by CRL1^BTRC^ or CRL1^FBXW11^ (also referred to as CRL1^βTRCP1^ and CRL1^βTRCP2^)^55^ (Fig. 5g). Accordingly, these SBMs and substrates were selectively identified by our active CRL probing method upon cell treatment with TNF (Fig. 5f, Extended data Fig. 5e). Remarkably, it also identified other key components of the TNF-regulated degradation and signaling pathways: specifically, the kinase CHUK responsible for generating the NFKBIA and NFKBIE phospho-degrons, and the transcription factors NFKB1 and RELA (Fig. 5f). Thus, our workflow also identifies proteins involved in signaling pathways associated with ubiquitylated substrates.

### Baseline active CRL repertoire primes cellular response

We next addressed the fundamental question if active CRL repertoires vary in different cell types by quantitatively comparing cellular repertoires of neddylated CRLs without endogenous tagging of cullins. Thus, we used our proteomics pipeline to probe neddylated CRL repertoires across a panel of ten cell lines, derived from kidney, tongue, brain, blood, bone, lung, ovary and prostate. To normalize for intrinsic differences (Extended data Fig. 6), we compared relative loss of SBMs in N8C_Fab3b IPs arising from 2-hour MLN4924 treatment. Remarkably, the relative levels of 83 SBMs changed significantly in at least one cell line (Fig. 6a). Several SBMs, for example, BTRC (aka βTRCP1), KLHL12 and CRBN, show large variations in neddylated CRL occupancies in different lines, while VHL was highly associated except in CAL33 cells (Fig. 6b-d).

**Fig. 6:**
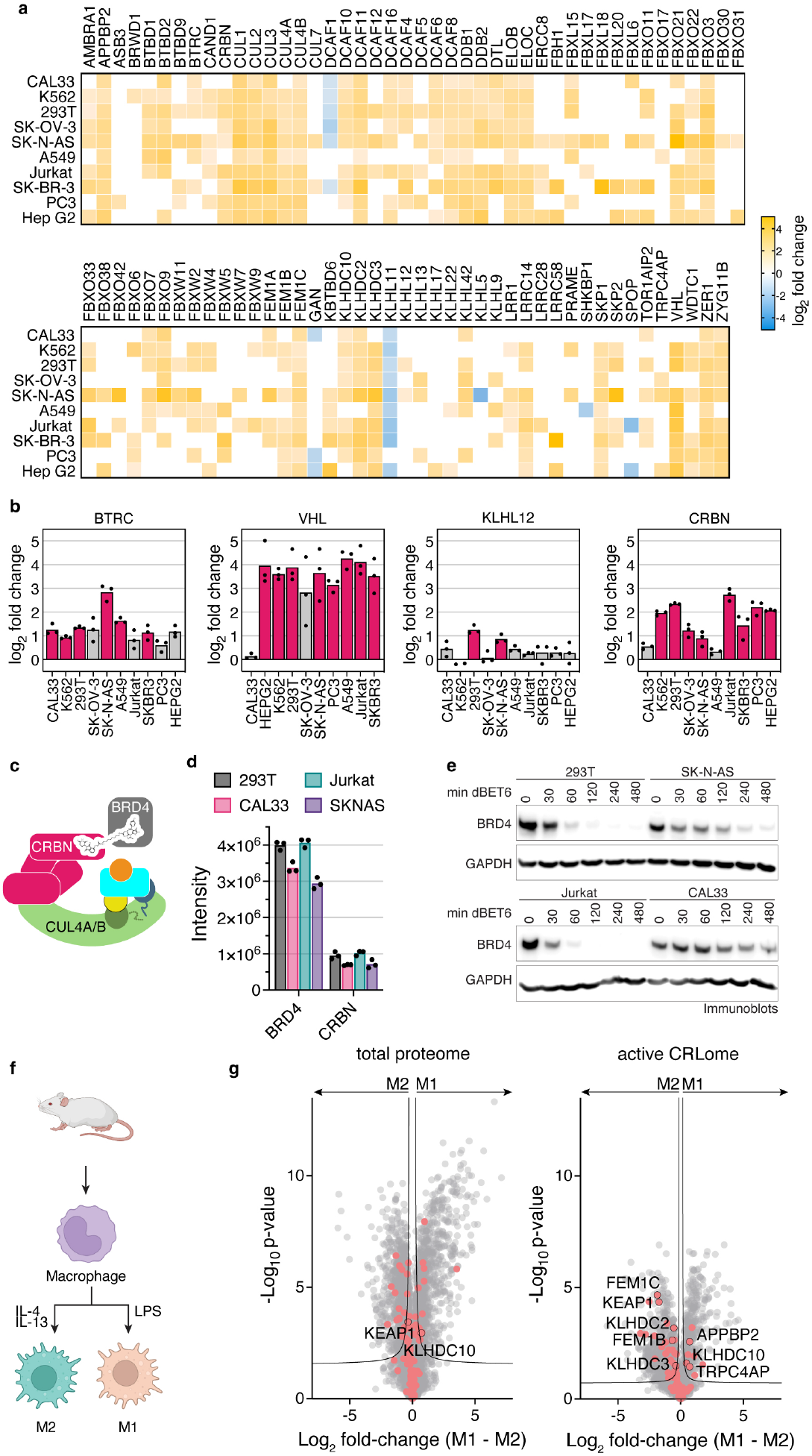
Baseline active CRL repertoire primes cellular response. **a**, Heatmap representation of N8C_Fab3b based CRL profiling in different human cell lines. Log_2_ fold change comparing DMSO to MLN4924 treated cells with all listed SBMs being significantly enriched over control in at least one cell line. SBMs not significantly enriched over control in a particular cell line are not colored in. **b**, Bar graphs based on (a), representing differences in the identified levels of SBMs BTRC, VHL, KLH12 and CRBN. Cell lines with pink bars show a significant difference in their N8C_Fab3b profiling results after DMSO or MLN4924 treatment. **c**, Cartoon representation of the NEDD8-CUL4^CRBN^ complex hijacked by dBET6 for degradation of BRD4. **d**, Bar graph of protein group intensities as determined by total proteomics as a readout of cellular protein levels for BRD4 and CRBN in 293T, CAL33, Jurkat, SKNAS. **e**, Time course of BRD4 degradation induced by treatment with 0.1 µM dBET6 of 293T, SK-N-AS, Jurkat and CAL33 for indicated durations. BRD4 degradation was monitored by immunoblotting using GAPDH as a loading control. **f**, Schematic of production of mouse bone marrow derived macrophages and activation to M1 and M2 macrophages by treatment with LPS or a combination of IL-4 and IL-13, respectively. Created with BioRender.com. **g**, Volcano plots of the differences between M1 and M2 mouse macrophages seen in the total proteome (left) and active CRLome as determined by N8C_Fab3b based profiling (right). Known CRL components are indicated in red and proteins of interest labeled with their gene names. Curve for 5% false discovery rate (s0 = 0.1) is shown.

Can pre-assembly into an active CRL impact response to a degradation signal? We tested this by examining dBET6-induced degradation of BRD4, mediated by neddylated CRL4^CRBN56^(Fig. 6c). Four cell lines were selected spanning the range of CRBN assembly with neddylated cullins, high in 293T and Jurkat cells, lower in SK-N-AS and very low in CAL33. CRBN levels in N8C_Fab3b IPs largely correlated with its expression. However, SK-N-AS and CAL33 cells showed nearly identical CRBN quantities but different degrees of assembly into active CRL complexes, indicating that formation of neddylated CRL4^CRBN^ also depends on factors beyond protein levels (Fig. 6d). Degradation efficiency correlated most strongly with the degree of CRBN assembly into a neddylated CRL as determined by our probe (Fig. 6e).

Conservation of the cullin WHB domain and NEDD8 sequences suggested that our profiling method could be extended to cells from other mammals (Extended Data Fig. 7a and b). Indeed, N8C_Fab3b IPs also enriched murine cullins, adapter proteins and SBMs in a neddylation-dependent manner (i.e. MLN4924-sensitive, Extended Data Fig. 8a), which afforded the opportunity to probe active CRL repertoires from cells derived from a primary source (Extended Data Fig. 8b). We therefore investigated differences across macrophage activation, a robust ex vivo model of cell state changes related to anti-microbial (“M1”) or anti-helminth and tissue reparative (“M2”) functions^57,58^. Mouse macrophages were differentiated from bone marrow progenitors with CSF1 and left unstimulated (“M0”), or stimulated with LPS or IL-4 plus IL-13 to form M1 or M2 activation states, respectively (Fig. 6f, Extended Data Fig. 8c). Total proteome analyses showed that M1 and M2 macrophages have generally similar levels of CRL components. However, probing with N8C_Fab3b revealed considerable differences in their neddylated CRL repertoires (Fig. 6g). Interestingly, the 26 potential SBMs differing between the M1 and M2 states include seven of the eight known to bind “C-degrons”, which are specific sequences at protein C-termini^59,60^. C-degrons are thought to be generated in stress conditions that trigger mis-translation or proteolytic cleavage. Notably, the redox sensing SBMs KEAP1^61,62^ and FEM1B^63,64^ also stand out as maintained between M0 and M2, but relatively decreased amongst the CRLs active in M1, consistent with their distinct metabolism and roles in inflammatory responses^57,58,65^. Our data imply overall that distinct CRLs are activated during quality control pathways differentially deployed by M1 or M2 states.

## Discussion

In this work, we started by asking if it was possible to generate affinity reagent(s) selectively binding a site-specifically UBL-modified target. We devised a negative>negative>positive strategy for selecting binders amongst phage displayed Fabs, applied this to two distinct neddylated cullins, and performed subsequent affinity maturation. Our strategy yielded eight affinity reagents selectively binding neddylated cullins in solution and in immunoblots, six of which specifically IP neddylated cullins and protect CUL1 and/or CUL2 from CSN-mediated deneddylation (Figs. 1-2). Upon publication we will provide the reagent collection to the community, while here deeply characterizing and developing one - N8C_Fab3b, which recovers neddylated CUL1, CUL2, CUL3, CUL4 and their associated proteins in IPs (Fig. 4d).

Remarkably, N8C_Fab3b binds a biologically-relevant conformation: the specific arrangement of NEDD8-linked to CUL1 in active CRL1 E3s (Fig. 3). While this conformation has so far only been structurally visualized for CUL1-based complexes with ubiquitin-carrying enzymes^11,12^, the binding of N8C_Fab3b provides biochemical evidence that this active conformation is conserved for NEDD8 linked to cullins 1-4. Remarkably, this Fab leaves NEDD8’s I44-patch free to bind ARIH1 during ubiquitylation (Fig. 3d, Extended Data Fig. 2f). Unlike E2 enzymes, which readily disengage from neddylated CRL catalytic domains^66^, ARIH1 binds even when not performing ubiquitylation^24^. Accordingly, ARIH1 copurifies with numerous CRLs in a neddylation-dependent manner, and plays crucial roles in NEDD8-dependent CRL functions^24,67^. Thus, the capacity to accommodate ARIH1 could be important for surveying the cellular landscapes of neddylated CRLs, and our mechanistic data showed that N8C_Fab3b could achieve this feat.

Performing IPs with N8C_Fab3b in various settings revealed fundamental features of the neddylated CRL network. For example, we discovered that there is not a singular effect of inhibiting neddylation or deneddylation on SBM association across the system of neddylated cullins (Fig. 4d). For a small subset of SBMs queried in our initial survey, association with neddylated cullins is impaired by both MLN4924 and CSNi5. This includes CRBN, which undergoes autodegradation in the absence of substrate(Mayor-Ruiz et al., 2019). Thus, loss of CRBN and such autoregulated SBMs would be expected from MLN4924 eliminating neddylation, and CSNi5 promoting degradation. However, the levels of most SBMs detected in N8C_Fab3b IPs decrease upon treatment with MLN4924 but not CSNi5. Considering that prior studies showed that the repertoire of SBMs co-IP’ing with endogenously tagged CUL1 or CUL4 shifts upon cell treatment with a variety of extracellular stimuli^40,42^, this suggested N8C_Fab3b IPs could be used to identify pathways stimulated by signals but without requiring laborious endogenous tagging.

We developed a robust and portable proteomics pipeline employing N8C_Fab3b that identifies the SBMs - and in some cases their substrates - responding to signals. Moreover, for the TNF stimulus, the pipeline not only identified E3s and their substrates, but also the kinase producing the phosphodegrons required to bind the identified SBMs, and components of the transcriptional complex liberated by the neddylated CRL pathway (Fig. 5). Although the underlying protein interaction architectures must await future studies, the data clearly show how our workflow can illuminate entire signaling pathways. Interestingly, another neddylated CRL1 complex, with the SBM SKP2, forms a higher-order assembly that includes its substrate p27 and Cyclin-CDK2-CKSHS1 kinases that produce the phosphodegron^12,20^. Thus, our results suggest that such kinase-substrate-E3 ligase signaling complexes - potentially formed transiently - may be more widespread than previously appreciated.

A key feature of our pipeline is that it can be applied in a generic manner in any system. We found striking variation of more than 70 SBMs across different cell lines (Fig. 6a). Pursuing targeted protein degradation mediated by one of them (CRBN) showed there is an overall link between its protein levels and degradation efficiency. However, targeted protein degradation efficiency correlated better with CRBN association with neddylated cullins. Furthermore, we asked if we could apply the probe to gain new insights into a system not readily amenable to endogenous tagging, originating from a mouse. We discovered that neddylated CRL repertoires vary across macrophage activation states, with noteworthy differences in E3s associated with quality control and stress responses (Fig. 6g). Amongst the variations are levels of C-degron binding SBMs, including KLHDC10 associated with ribosome quality control, and KLHDC2 that has recently gained prominence as a potential handle for targeted protein degradation^68,69^. Thus, not only do CRL networks dynamically rearrange to drive cellular signaling, but we propose that CRL repertoires may also adjust to resolve stresses arising from toxic effectors such as those required for diverse macrophage activities that include microbial killing, efferocytosis and tissue repair^57,58,65^.

Finally, this study highlights the potential of harnessing PTMs to probe active complexes mediating dynamic cellular signaling. Utilizing binders to recognize an activating PTM, or a conformation unique to active complexes, provides an alternative approach to select active complexes from amongst the broader cellular pool of constituent molecules. While the ubiquitin system has classically been probed by small-molecules reacting with enzyme catalytic cysteines, such as deubiquitinating enzymes and E3 ligases in the HECT, RBR, and RCR families^70-74^, we show utility of conformation-specific affinity probes to ensnare E3 complexes lacking residues easily targeted by reactive chemical moieties. Our approach selectively targeting a site-specific modification - and in so doing capturing activated forms of numerous CRL complexes with multiple cullin backbones, from across mammalian species, without prior engineering of the cellular system of interest and with a generally applicable proteomic platform - enables new insights into dynamic E3 ligase and targeted protein degradation pathways.

## Acknowledgments

We thank all the members of our labs for their input and support. In particular we thank D. Scott, D. Miller, S. Bozeman, J. Kellermann, J.R. Prabu, I. Paron and T. Heymann for assistance, reagents and helpful discussions. This work was funded by the ERC under the H2020 research and innovation programme (ADG-789016-Nedd8Activate), by the DFG (SCHU3196/1-1, Gottfried Wilhelm Leibniz Prize), ALSAC, St. Jude Children’s Research Hospital, NIH P30CA021765 (to BAS); the Max Planck Society (to MM and BAS); and Canadian Institutes of Health Research grant MOP-93684 (to SSS).

## Author Contributions

LTH, JS, DMD, SSS, BAS conceived the project. LTH, DMD, KB. DY generated proteins. JS performed Fab selections, ELISAs and initial characterization. LTH, DMD, KB, DY performed additional biochemical characterization of the Fab suite and biochemical assays. DMD obtained crystal structure. LTH performed cellular assays, developed the active CRL profiling pipeline, and acquired and analyzed mass spectrometry data. PJM generated and advised on experiments with BMDMs. MM, SSS, BAS supervised the project. LTH and BAS wrote the manuscript, with input from all authors.

## Data availability

This is a presubmission preprint. Extended data figures, methods, Fab sequences and expression constructs, crystallographic and proteomics data will be available upon publication in a journal.

## References

1 Komander, D. & Rape, M. The ubiquitin code. Annual Review of Biochemistry 81, 203–229 (2012). https://doi.org:10.1146/annurev-biochem-060310-170328

2 Husnjak, K. & Dikic, I. Ubiquitin-binding proteins: decoders of ubiquitin-mediated cellular functions. Annu Rev Biochem 81, 291–322 (2012). https://doi.org:10.1146/annurev-biochem-051810-094654

3 Matsumoto, M. L. et al. K11-Linked Polyubiquitination in Cell Cycle Control Revealed by a K11 Linkage-Specific Antibody. Mol Cell 39, 477–484 (2010). https://doi.org:10.1016/j.molcel.2010.07.001

4 Yau, R. G. et al. Assembly and Function of Heterotypic Ubiquitin Chains in Cell-Cycle and Protein Quality Control. Cell 171, 918-933.e920 (2017). https://doi.org:10.1016/j.cell.2017.09.040

5 Michel, M. A., Swatek, K. N., Hospenthal, M. K. & Komander, D. Ubiquitin Linkage-Specific Affimers Reveal Insights into K6-Linked Ubiquitin Signaling. Mol Cell 68, 233-246.e235 (2017). https://doi.org:10.1016/j.molcel.2017.08.020

6 Yu, Y. et al. K29-linked ubiquitin signaling regulates proteotoxic stress response and cell cycle. Nat Chem Biol 17, 896–905 (2021). https://doi.org:10.1038/s41589-021-00823-5

7 Sims, J. J. & Cohen, R. E. Linkage-specific avidity defines the lysine 63-linked polyubiquitin-binding preference of rap80. Mol Cell 33, 775–783 (2009). https://doi.org:10.1016/j.molcel.2009.02.011

8 Armstrong, A. A., Mohideen, F. & Lima, C. D. Recognition of SUMO-modified PCNA requires tandem receptor motifs in Srs2. Nature 483, 59–63 (2012). https://doi.org:10.1038/nature10883

9 Panier, S. et al. Tandem protein interaction modules organize the ubiquitin-dependent response to DNA double-strand breaks. Mol Cell 47, 383–395 (2012). https://doi.org:10.1016/j.molcel.2012.05.045

10 Morgan, M. T. & Wolberger, C. Recognition of ubiquitinated nucleosomes. Curr Opin Struct Biol 42, 75–82 (2017). https://doi.org:10.1016/j.sbi.2016.11.016

11 Baek, K. et al. NEDD8 nucleates a multivalent cullin–RING–UBE2D ubiquitin ligation assembly. Nature 578, 461–466 (2020). https://doi.org:10.1038/s41586-020-2000-y

12 Horn-Ghetko, D. et al. Ubiquitin ligation to F-box protein targets by SCF-RBR E3-E3 super-assembly. Nature 590 (2021). https://doi.org:10.1038/s41586-021-03197-9

13 Pan, Z. Q., Kentsis, A., Dias, D. C., Yamoah, K. & Wu, K. Nedd8 on cullin: building an expressway to protein destruction. Oncogene 23, 1985–1997 (2004).

14 Vogl, A. M. et al. Global site-specific neddylation profiling reveals that NEDDylated cofilin regulates actin dynamics. Nat Struct Mol Biol 27 (2020). https://doi.org:10.1038/s41594-019-0370-3

15 Lobato-Gil, S. et al. Proteome-wide identification of NEDD8 modification sites reveals distinct proteomes for canonical and atypical NEDDylation. Cell Rep 34, 108635 (2021). https://doi.org:10.1016/j.celrep.2020.108635

16 Willems, A. R., Schwab, M. & Tyers, M. A hitchhiker’s guide to the cullin ubiquitin ligases: SCF and its kin. Biochimica et biophysica acta 1695, 133–170 (2004).

17 Rusnac, D. V. & Zheng, N. Structural Biology of CRL Ubiquitin Ligases. Adv Exp Med Biol 1217, 9–31 (2020). https://doi.org:10.1007/978-981-15-1025-0_2

18 Wang, K., Deshaies, R. J. & Liu, X. Assembly and Regulation of CRL Ubiquitin Ligases. Adv Exp Med Biol 1217, 33–46 (2020). https://doi.org:10.1007/978-981-15-1025-0_3

19 Harper, J. W. & Schulman, B. A. Cullin-RING Ubiquitin Ligase Regulatory Circuits: A Quarter Century Beyond the F-Box Hypothesis. Annual Review of Biochemistry 90, 403–429 (2021). https://doi.org:10.1146/annurev-biochem-090120-013613

20 Skaar, J. R., Pagan, J. K. & Pagano, M. Mechanisms and function of substrate recruitment by F-box proteins. Nature reviews. Molecular cell biology 14, 369–381 (2013). https://doi.org:10.1038/nrm3582

21 Duda, D. M. et al. Structural insights into NEDD8 activation of cullin-RING ligases: conformational control of conjugation. Cell 134, 995–1006 (2008). https://doi.org:10.1016/j.cell.2008.07.022

22 Yamoah, K. et al. Autoinhibitory regulation of SCF-mediated ubiquitination by human cullin 1’s C-terminal tail. P Natl Acad Sci USA 105, 12230–12235 (2008). https://doi.org:10.1073/pnas.0806155105

23 Saha, A. & Deshaies, R. J. Multimodal Activation of the Ubiquitin Ligase SCF by Nedd8 Conjugation. Mol Cell 32, 21–31 (2008). https://doi.org:10.1016/j.molcel.2008.08.021

24 Scott, D. C. et al. Two Distinct Types of E3 Ligases Work in Unison to Regulate Substrate Ubiquitylation. Cell 166, 1198-+ (2016). https://doi.org:10.1016/j.cell.2016.07.027

25 Baek, K., Scott, D. C. & Schulman, B. A. NEDD8 and ubiquitin ligation by cullin-RING E3 ligases. Curr Opin Struc Biol 67, 101–109 (2021). https://doi.org:10.1016/j.sbi.2020.10.007

26 Angers, S. et al. Molecular architecture and assembly of the DDB1-CUL4A ubiquitin ligase machinery. Nature 443, 590–593 (2006). https://doi.org:10.1038/nature05175

27 Cardote, T. A. F., Gadd, M. S. & Ciulli, A. Crystal Structure of the Cul2-Rbx1-EloBC-VHL Ubiquitin Ligase Complex. Structure 25, 901–911 e903 (2017). https://doi.org:10.1016/j.str.2017.04.009

28 Kostrhon, S. et al. CUL5-ARIH2 E3-E3 ubiquitin ligase structure reveals cullin-specific NEDD8 activation. Nat Chem Biol 17, 1075–1083 (2021). https://doi.org:10.1038/s41589-021-00858-8

29 Fischer, Eric S. et al. The Molecular Basis of CRL4DDB2/CSA Ubiquitin Ligase Architecture, Targeting, and Activation. Cell 147, 1024–1039 (2011). https://doi.org:https://doi.org/10.1016/j.cell.2011.10.035

30 Emberley, E. D., Mosadeghi, R. & Deshaies, R. J. Deconjugation of Nedd8 from Cul1 is directly regulated by Skp1-F-box and substrate, and the COP9 signalosome inhibits deneddylated SCF by a noncatalytic mechanism. J Biol Chem 287, 29679–29689 (2012). https://doi.org:10.1074/jbc.M112.352484

31 Enchev, Radoslav I. et al. Structural Basis for a Reciprocal Regulation between SCF and CSN. Cell Reports 2, 616–627 (2012). https://doi.org:https://doi.org/10.1016/j.celrep.2012.08.019

32 Lingaraju, G. M. et al. Crystal structure of the human COP9 signalosome. Nature 512, 161–165 (2014). https://doi.org:10.1038/nature13566

33 Cavadini, S. et al. Cullin-RING ubiquitin E3 ligase regulation by the COP9 signalosome. Nature 531, 598–603 (2016). https://doi.org:10.1038/nature17416

34 Mosadeghi, R. et al. Structural and kinetic analysis of the COP9-Signalosome activation and the cullin-RING ubiquitin ligase deneddylation cycle. Elife 5 (2016). https://doi.org:10.7554/eLife.12102

35 Mayor-Ruiz, C. et al. Plasticity of the Cullin-RING Ligase Repertoire Shapes Sensitivity to Ligand-Induced Protein Degradation. Mol Cell 75, 849-858.e848 (2019). https://doi.org:https://doi.org/10.1016/j.molcel.2019.07.013

36 Mayor-Ruiz, C. et al. Rational discovery of molecular glue degraders via scalable chemical profiling. Nat Chem Biol 16, 1199–1207 (2020). https://doi.org:10.1038/s41589-020-0594-x

37 Wu, T. et al. Targeted protein degradation as a powerful research tool in basic biology and drug target discovery. Nat Struct Mol Biol 27, 605–614 (2020). https://doi.org:10.1038/s41594-020-0438-0

38 Dale, B. et al. Advancing targeted protein degradation for cancer therapy. Nature Reviews Cancer 21, 638–654 (2021). https://doi.org:10.1038/s41568-021-00365-x

39 Bennett, E. J., Rush, J., Gygi, S. P. & Harper, J. W. Dynamics of cullin-RING ubiquitin ligase network revealed by systematic quantitative proteomics. Cell 143, 951–965 (2010). https://doi.org:10.1016/j.cell.2010.11.017

40 Reitsma, J. M. et al. Composition and Regulation of the Cellular Repertoire of SCF Ubiquitin Ligases. Cell 171, 1326-1339.e1314 (2017). https://doi.org:https://doi.org/10.1016/j.cell.2017.10.016

41 Liu, X. et al. Cand1-Mediated Adaptive Exchange Mechanism Enables Variation in F-Box Protein Expression. Mol Cell 69, 773-786.e776 (2018). https://doi.org:10.1016/j.molcel.2018.01.038

42 Reichermeier, K. M. et al. PIKES Analysis Reveals Response to Degraders and Key Regulatory Mechanisms of the CRL4 Network. Mol Cell 77, 1092-1106.e1099 (2020). https://doi.org:10.1016/j.molcel.2019.12.013

43 Adams, J. J. & Sidhu, S. S. Synthetic antibody technologies. Curr Opin Struc Biol 24, 1–9 (2014). https://doi.org:https://doi.org/10.1016/j.sbi.2013.11.003

44 Miersch, S. et al. Scalable High Throughput Selection From Phage-displayed Synthetic Antibody Libraries. Jove-J Vis Exp (2015). https://doi.org:ARTNe5149210.3791/51492

45 Hornsby, M. et al. A High Through-put Platform for Recombinant Antibodies to Folded Proteins. Molecular & Cellular Proteomics 14, 2833–2847 (2015). https://doi.org:10.1074/mcp.O115.052209

46 Soucy, T. A. et al. An inhibitor of NEDD8-activating enzyme as a new approach to treat cancer. Nature 458, 732–736 (2009). https://doi.org:10.1038/nature07884

47 Schlierf, A. et al. Targeted inhibition of the COP9 signalosome for treatment of cancer. Nat Commun 7, 13166–13166 (2016). https://doi.org:10.1038/ncomms13166

48 Han, T. et al. Anticancer sulfonamides target splicing by inducing RBM39 degradation via recruitment to DCAF15. Science 356, eaal3755–eaal3755 (2017). https://doi.org:10.1126/science.aal3755

49 Uehara, T. et al. Selective degradation of splicing factor CAPERα By anticancer sulfonamides. Nat Chem Biol 13, 675–680 (2017). https://doi.org:10.1038/nchembio.2363

50 Zengerle, M., Chan, K.-H. & Ciulli, A. Selective Small Molecule Induced Degradation of the BET Bromodomain Protein BRD4. ACS Chemical Biology 10, 1770–1777 (2015). https://doi.org:10.1021/acschembio.5b00216

51 Gadd, M. S. et al. Structural basis of PROTAC cooperative recognition for selective protein degradation. Nat Chem Biol 13, 514–521 (2017). https://doi.org:10.1038/nchembio.2329

52 Cowan, A. D. & Ciulli, A. Driving E3 Ligase Substrate Specificity for Targeted Protein Degradation: Lessons from Nature and the Laboratory. Annu Rev Biochem 91, 295–319 (2022). https://doi.org:10.1146/annurev-biochem-032620-104421

53 Vashisht, A. A. et al. Control of iron homeostasis by an iron-regulated ubiquitin ligase. Science 326, 718–721 (2009). https://doi.org:10.1126/science.1176333

54 Salahudeen, A. A. et al. An E3 Ligase Possessing an Iron-Responsive Hemerythrin Domain Is a Regulator of Iron Homeostasis. Science 326, 722–726 (2009). https://doi.org:10.1126/science.1176326

55 Kanarek, N., London, N., Schueler-Furman, O. & Ben-Neriah, Y. Ubiquitination and degradation of the inhibitors of NF-kappaB. Cold Spring Harb Perspect Biol 2, a000166 (2010). https://doi.org:10.1101/cshperspect.a000166

56 Winter, G. E. et al. BET Bromodomain Proteins Function as Master Transcription Elongation Factors Independent of CDK9 Recruitment. Mol Cell 67, 5-+ (2017). https://doi.org:10.1016/j.molcel.2017.06.004

57 Murray, P. J. et al. Macrophage activation and polarization: nomenclature and experimental guidelines. Immunity 41, 14–20 (2014). https://doi.org:10.1016/j.immuni.2014.06.008

58 Murray, P. J. Macrophage Polarization. Annu Rev Physiol 79, 541–566 (2017). https://doi.org:10.1146/annurev-physiol-022516-034339

59 Koren, I. et al. The Eukaryotic Proteome Is Shaped by E3 Ubiquitin Ligases Targeting C-Terminal Degrons. Cell 173, 1622-1635.e1614 (2018). https://doi.org:https://doi.org/10.1016/j.cell.2018.04.028

60 Lin, H. C. et al. C-Terminal End-Directed Protein Elimination by CRL2 Ubiquitin Ligases. Mol Cell 70, 602–613 e603 (2018). https://doi.org:10.1016/j.molcel.2018.04.006

61 Ishii, T. et al. Transcription factor Nrf2 coordinately regulates a group of oxidative stress-inducible genes in macrophages. Journal of Biological Chemistry 275, 16023–16029 (2000). https://doi.org:DOI10.1074/jbc.275.21.16023

62 Dinkova-Kostova, A. T., Kostov, R. V. & Canning, P. Keap1, the cysteine-based mammalian intracellular sensor for electrophiles and oxidants. Arch Biochem Biophys 617, 84–93 (2017). https://doi.org:10.1016/j.abb.2016.08.005

63 Manford, A. G. et al. A Cellular Mechanism to Detect and Alleviate Reductive Stress. Cell 183, 46–61 e21 (2020). https://doi.org:10.1016/j.cell.2020.08.034

64 Manford, A. G. et al. Structural basis and regulation of the reductive stress response. Cell 184, 5375–5390 e5316 (2021). https://doi.org:10.1016/j.cell.2021.09.002

65 Bakker, N. V. & Pearce, E. J. Cell-intrinsic metabolic regulation of mononuclear phagocyte activation: Findings from the tip of the iceberg. Immunological Reviews 295, 54–67 (2020). https://doi.org:10.1111/imr.12848

66 Eletr, Z. M., Huang, D. T., Duda, D. M., Schulman, B. A. & Kuhlman, B. E2 conjugating enzymes must disengage from their E1 enzymes before E3-dependent ubiquitin and ubiquitin-like transfer. Nat Struct Mol Biol 12, 933–934 (2005).

67 Zhang, Y. et al. Adaptive exchange sustains cullin-RING ubiquitin ligase networks and proper licensing of DNA replication. Proc Natl Acad Sci U S A 119, e2205608119 (2022). https://doi.org:10.1073/pnas.2205608119

68 Schapira, M., Calabrese, M. F., Bullock, A. N. & Crews, C. M. Targeted protein degradation: expanding the toolbox. Nat Rev Drug Discov 18, 949–963 (2019). https://doi.org:10.1038/s41573-019-0047-y

69 Thrun, A. et al. Convergence of mammalian RQC and C-end rule proteolytic pathways via alanine tailing. Mol Cell 81, 2112–2122 e2117 (2021). https://doi.org:10.1016/j.molcel.2021.03.004

70 Pao, K. C. et al. Probes of ubiquitin E3 ligases enable systematic dissection of parkin activation. Nat Chem Biol 12, 324–331 (2016). https://doi.org:10.1038/nchembio.2045

71 Mulder, M. P. C., Witting, K. F. & Ovaa, H. Cracking the Ubiquitin Code: The Ubiquitin Toolbox. Curr Issues Mol Biol 37, 1–20 (2020). https://doi.org:10.21775/cimb.037.001

72 Henneberg, L. T. & Schulman, B. A. Decoding the messaging of the ubiquitin system using chemical and protein probes. Cell Chem Biol 28, 889–902 (2021). https://doi.org:10.1016/j.chembiol.2021.03.009

73 Zheng, Q., Su, Z., Yu, Y. & Liu, L. Recent progress in dissecting ubiquitin signals with chemical biology tools. Curr Opin Chem Biol 70, 102187 (2022). https://doi.org:10.1016/j.cbpa.2022.102187

74 Pao, K. C. et al. Activity-based E3 ligase profiling uncovers an E3 ligase with esterification activity. Nature 556, 381–385 (2018). https://doi.org:10.1038/s41586-018-0026-1

